# Infrared gas analysis as a method of measuring seagrass photosynthetic rate in the face of desiccation stress

**DOI:** 10.1101/2024.01.16.575902

**Authors:** Kyle A Capistrant-Fossa, Kenneth H Dunton

## Abstract

Photosynthesis, a core autotrophic metabolic process for aquatic and terrestrial organisms, is the backbone of the global carbon biogeochemical cycle. Inorganic assimilation of carbon in photosynthesis is relative difficult to measure in an aqueous medium since carbon readily reacts with ions in water. Therefore, aquatic photosynthesis is often measured using secondary methods that introduce uncertainty into measurements (e.g., oxygen evolution). One technique, infrared gas analysis (IRGA), uses a closed gas loop to calculate an accurate carbon budget. Multiple studies have successfully used IRGA with intertidal seagrasses, but it remains unknown how applicable the technology is for underwater plants. Here, we evaluate the potential of IRGA to measure carbon assimilation of subtidal seagrasses temporarily removed from seawater, and evaluate how carbon fixation rates and chlorophyll fluorescence characteristics of subtidal seagrasses change as they desiccate. We use IRGA for four common seagrass species from the Western Gulf of Mexico (*Halophila engelmannii, Halodule wrightii, Syringodium filiforme*, and *Thalassia testudinum*) paired with pulse amplitude modulated fluorometry to measure desiccation stress. *Halophila* had the highest maximum carbon assimilation rate (6.06 *µ*mol C m^−2^s^−1^), followed by *Thalassia* (5.58 *µ*mol C m^−2^ s^−1^), *Halodule* (4.75 *µ*mol C m^−2^ s^−1^), and *Syringodium* (3.63 *µ*mol C m^−2^ s^−1^). *Thalassia* was most resistant to desiccation stress as reflected by the plant’s ability to maintain high maximum leaf quantum efficiency (Fv/Fm) while the other species were not. Additionally, *Thalassia* had a slower desiccation rate (2.3% min^−1^ cm^−2^) than 4.79% *Syringodium filiforme* (4.79% min^−1^ cm^−2^) and *Halodule wrightii* (30.17% min^−1^ cm^−2^). Together, our provide reasonable measures of carbon assimilation and support previous studies of seagrass desiccation stress gradients along depth. Overall, we recognize IRGA as a promising direction for future studies of seagrass productivity and recommend further investigation.

## INTRODUCTION

Photosynthesis, a key autotrophic metabolic process, allows chlorophyll-containing organisms to utilize solar electromagnetic radiation to convert inorganic carbon dioxide and water into organic polysaccharides and oxygen. Ultimately, this process fuels diverse terrestrial and aquatic ecosystems through supporting heterotrophic metabolism by providing oxygen for cellular respiration and facilitating nutrient cycling. Photosynthetic autotrophy is a successful evolutionary niche leading to the rise of about *∼*72,000 algae species (primarily aquatic; Guiry, 2012) and *∼*374,000 plant species (primarily terrestrial; Christenhusz and Byng, 2016). Globally, these organisms are the backbone of the carbon biogeochemical cycle and fix 120 – 175 gross Pg yr^−1^ of carbon with aquatic and terrestrial organisms each contributing roughly half (Welp et al., 2011; Falkowski and Raven, 2013). These estimates are largely based on countless in-situ measurements and experimental work that has provided insight into fundamental controls of plant photosynthesis.

In-situ productivity measurements have three common targets: carbon, oxygen, or chlorophyll fluorescence. Carbon fixation is the goal of plant photosynthesis; therefore, measuring carbon is ideal, but difficult to measure in an aquatic environment compared to terrestrial systems because 1) the large dissolved inorganic carbon (DIC) pools in aquatic environments create the need for costly and sensitive sensors, and 2) the complex carbonate equilibrium system requires measurements of pH, alkalinity, total DIC, and/or pCO_2_ (Silva et al., 2009). Radioactively tagged isotopes are commonly used to measure carbon assimilation in the field (i.e., ^14^C; Nielsen, 1951), but multiple physiological processes and varied incubation times limit the interpretability and reliability of data (see review by Milligan et al., 2015).

Oxygen sensors are widely adopted because of their low cost and high precision (Silva et al., 2009). However, oxygen budgets are not directly representative of carbon fixation, therefore oxygen-based measures introduce errors into productivity studies. Non-specific oxygen consuming process (e.g., microbial respiration, geochemical reactions, gas ebullition) can interfere with measurements. Furthermore, the ratio of oxygen produced to carbon fixed (i.e., photosynthetic quotient) varies significantly between species and physiological conditions (e.g., 0.1 – 4.2) introducing large error and uncertainty (Trentman et al., 2023).

Plant productivity can also be inferred by measuring plant fluorescence (e.g., pulse amplitude modulated fluorometry “PAM”) from photosystem II in response to rapid pulses of light because it is directly proportional to photosynthetic rate (Ralph and Gademann, 2005). There are various experimental configurations that allow for determination of plant photosynthetic capacity, efficiency, and stress using the same technology (Ralph and Gademann, 2005). Numerous studies have used PAM to evaluate seagrass stress and photosynthetic rate (e.g., Beer and Björk, 2000). However, fluorescence only measures electron transfer rate (which is not always equivalent to O_2_ evolution rates) and cannot be used to estimate respiration rates (Beer and Björk, 2000).

Given the pros and cons of traditional techniques, there is need for development of a unified, accurate measure of aquatic productivity. Carbon-based measurements using infrared gas analysis (IRGA) are promising for adaptation to the aquatic environment. Terrestrial systems create a CO_2_ budget in a closed air loop, but modern adaptations can utilize water instead (Egle and Ernst, 1949; Beer et al., 2014). Only a few studies have employed this technology to directly measure carbon uptake in intertidal populations of aquatic plants (Leuschner and Rees, 1993; Pérez-Lloré ns and Niell, 1993; Leuschner et al., 1998; Silva et al., 2005) because of additional desiccation stress to plants.

Desiccation is a common stressor that affects aquatic plants and has been the focus of ecologists and biologists for decades (e.g., Stephenson and Stephenson, 1949). Typically, desiccation stress decreases the photosynthetic performance of a plant, but rates will usually rebound after rehydration (Dring, 1982). Globally, seaweeds will arrange themselves along shorelines based on differences in their desiccation tolerance (upper limit) and competitive ability (lower limit; Stephenson and Stephenson, 1949). Interestingly, seagrasses are an exception to this common pattern because subtidal plants are more desiccation tolerant compared to their shallow intertidal counterparts (Bjö rk et al., 1999; Shafer et al., 2007). Therefore, subtidal seagrasses are an excellent system to test the viability of infrared gas analysis to measure productivity in submerged aquatic plants because they may be tolerant of desiccation during measurements. Here we will utilize a terrestrial IRGA system to measure productivity rates and use a PAM fluorometer to understand plant stress levels. We hypothesize that short term emersion from seawater will not significantly stress plants and will allow for accurate productivity estimates.

## METHODS

### Seagrass Collection & Culture Methods

Whole plants, including intact roots and rhizomes, of *Thalassia testudinum* (Tt), *Halodule wrightii* (Hw), and *Halophila engelmannii* (He) were collected from Redfish Bay within the Mission Aransas National Estuarine Research Reserve and *Syringodium filiforme* (Sf) from East Flats within Corpus Christi Bay, Texas (Western Gulf of Mexico) in mid-May 2023. The seagrasses were washed with seawater to remove sediment and gently scraped with a razor blade to remove epiphytic organisms before being placed into one of three aquaria. Experimental specimens were cultured with gentle aeration at 20ºC, *∼*40 salinity, exposed to 50 *µ*mol photons m^−2^ s^−1^ photosynthetic proton flux density, and a 12:12 (Light:Dark) light cycle for 4 weeks. Blades were visually inspected before experiments and only those with good coloration, intact root systems, and low biofouling were used.

### Dead Seagrass Blades in LI-6400 Experiments

The commercial IRGA system used in this series of experiments, LI-6400 XT (LICOR, LINCOLN, NE), uses a dual water and carbon balance to measure productivity:

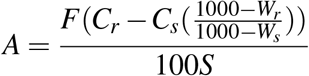

where A is assimilation rate *µ*mol CO_2_ m^−2^ s^−1^, F is air flow rate (*µ*mol s^−1^), C_*r*_ and C_*s*_ are sample and reference CO_2_ concentrations (*µ*mol CO_2_ (mol air)^−1^), and S is leaf area (cm^2^). However, W_*r*_ and W_*s*_ are molar dilution terms directly derived from transpiration, which can be assumed be 0 in seagrasses because seagrasses lack stomata, thereby reducing the formula to:

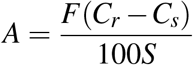

To determine the contribution of non-photosynthetic carbon assimilation (e.g., internal carbon accumulation) to net assimilation (i.e., A), we extracted all pigments from“dead” plants (n = 5/species) prior to experiments (see below) through repeated short exposures to a 1:1 95% Acetone 1N HCl solution followed by quenching in water until the plant leaf tissues were white.

### Seagrass Desiccation in LI-6400 Chamber

Seagrass blades were split into either“dead” or natural“experimental” treatments. Leaves (n = 5/species/treatment) were gently scraped of any epiphytes, wiped of excess water, weighed, trimmed to equal length, and placed into an LI-6400 XT standard leaf chamber equipped with a LED light source. The chamber was set to maintain a temperature of 20°C, the ambient humidity was maintained relatively constant between trials, the flow rate was set to between 250 – 500 *µ*mol s^−1^ depending on the desiccation rate of the blade, and photosynthetic proton flux density of 1000 *µ*mol photons m^−2^ s^−1^. Blades were removed from the chamber and weighed until they reached a constant weight (up to 30 minutes). General additive models were fit to describe the loss of water (i.e., relative biomass) as a function of time for each species and treatment. Differences between treatments were tested using generalized linear models assuming a gamma distribution. All data (DOI: 10.5281/zenodo.10475717) and code (https://github.com/kacf24/SeagrassDesiccation) is deposited online.

### Change in Carbon Fixation as Blades Desiccate

In parallel to the previous experiment (i.e., replicated experimental design) we measured carbon assimilation of seagrass blades measurements every 15 – 60 seconds for up to 30 minutes. General additive models were fit to describe carbon assimilation as a function of relative biomass for each species and treatment.

### Change in Fv/Fm and Rapid Light Curves

To evaluate our hypothesis that seagrasses will not be stressed by short emersions, we tracked changes in Fv/Fm (maximum leaf quantum efficiency), a measure of plant stress over time (Ralph and Gademann, 2005). We allowed dark-adapted seagrass blades (n = 5/species) to dry in from of a box fan for an hour. We measured Fv/Fm using a Walz Diving PAM and weighed the blades every 15 minutes. Additionally, we performed rapid light curves at the beginning and end of the 60 minutes with 10 second windows. Light curves were fit to the PGH model (Platt et al., 1980) and photosynethic parameters calculated using the phytotools package (Silsbe and Malkin, 2015).

### Timing and Frequency of Desiccation

To evaluate changes in desiccation across seasons, we approximate seagrass evapora-tion rates using a simple water evaporation model (Shi et al., 2017) that accounts for temperature, humidity, and windspeed:

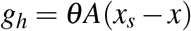

where *g*_*h*_ is the evaporation rate (kg H_2_O h^−1^), *θ* is the evaporation coefficient (kg m^−2^ h^−1^; empirically estimated using 25+ 19* air velocity), A is surface area (m^2^), x_*s*_ is the maximum humidity ratio of saturated air, and x is the humidity ratio of air. A was set to 1, and values of *θ*, x_*s*_, x were calculated using meteorological data obtained from a local airport (Mustang Beach, KRAS) from 1/1/2021 – 12/31/2022 and the psychrolib R Package (Jia, 2021). The major assumptions for this model are that 1) the ability of seagrasses to retain water does not significantly change over time and 2) water loss is primarily driven by abiotic factors. These assumptions are reasonable because seagrasses lack stomata used by other plants in biotic water regulation, they possess a thin, porous cuticle that is ineffective at water retention, and have a relatively low sulfated hygroscopic polysaccharide content compared to seaweeds (Den Hartog and Kuo, 2006). We used the mean low water datum (NOAA Tide Predictions, 8775237) as a reference with positive values representing inundation and negative representing exposure (Koch, 2001).

## RESULTS

### Seagrass Desiccation in LI-6400 Chamber

Seagrass biomass varied significantly among species and time (both p *<* 0.05) but not between dead and experimental treatments (p = 0.13; Table 1). *Syringodium* blades weighed the most on average (0.10 – 0.03 g cm^−2^), had the slowest rate of biomass loss (2.3% per min) and took 30 minutes to reach a constant biomass (Figure 1). In contrast, *Halophila* had the lowest biomass (0.01 – 0.004 g cm^−2^), the highest rate of biomass loss (12% per min) and reached a constant weight within 5 minutes (Figure 1). The biomass of *Thalassia* and *Halodule* blades overlapped with each other (0.03 – 0.01 g cm^−2^) and reached constant weight after 30 minutes of drying (Figure 1). However, *Halodule* lost biomass quicker (5.4% per min) and more variably than *Thalassia* (3.4% per min).

**Table 1.** Results of a generalized linear model of seagrass water loss over time between treated“dead” and natural“experimental” seagrass. The intercept is referenced to dead *Halophila engelmannii*.

| Biomass ∼ Treatment + Species + Time | |
| --- | --- |
| Term | p-value |
| Intercept | $10^{-16}$ |
| Species (Hw) | $10^{-16}$ |
| Species (Sf) | $10^{-16}$ |
| Species (Tt) | $10^{-16}$ |
| Treatment (Experimental) | 0.13 |
| Time | $10^{-16}$ |

**Figure 1.**
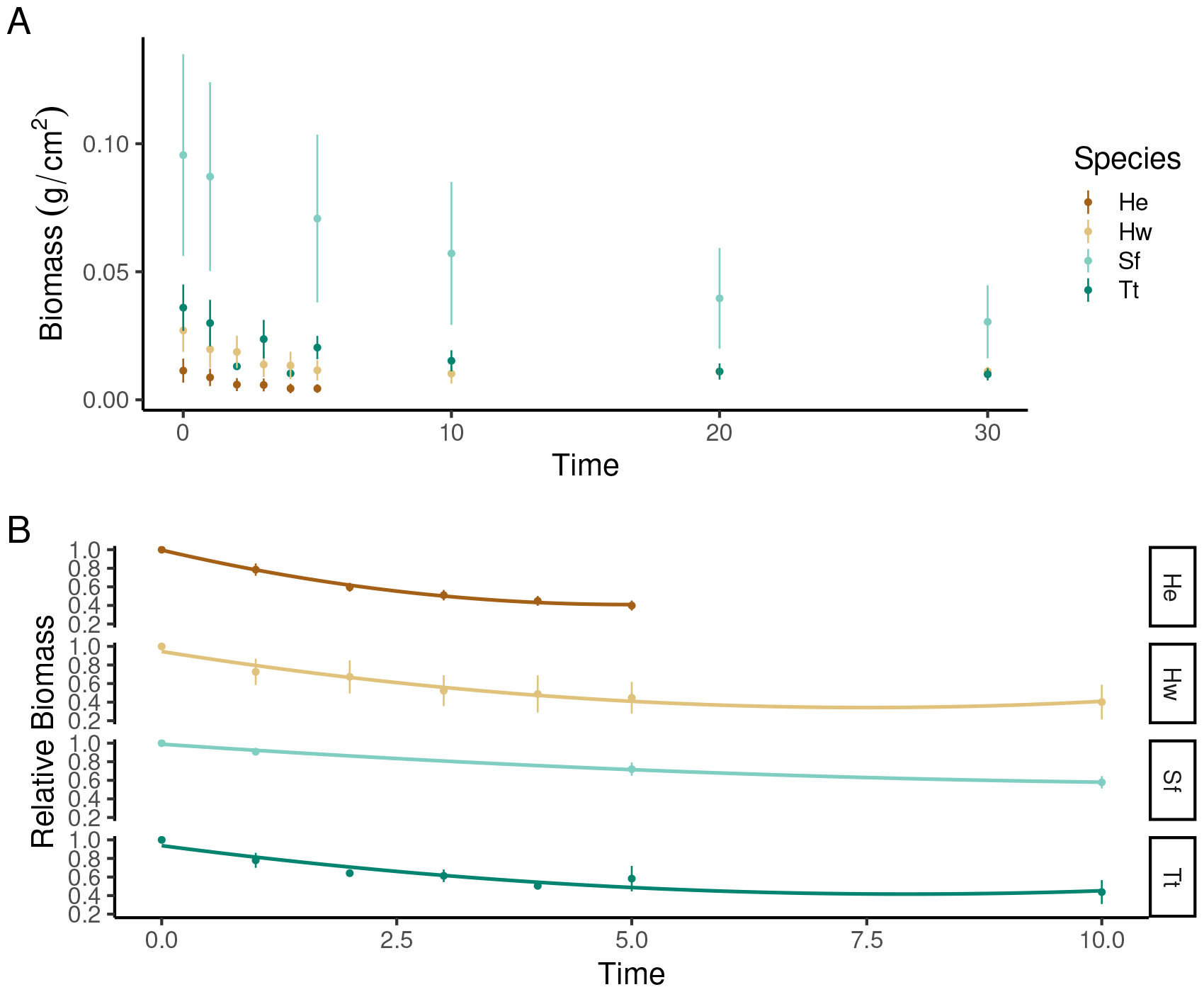
We found no significant (p *>* 0.05) differences between the biomass of dead and experimental seagrass blades, but the effects of time and species were significant (p*<* 0.05). Biomass is shown as both an absolute value (A) and a relative value (B) over time. Each point represents a mean± standard deviation (n = 5).

### Effects of Desiccation of Maximum Carbon Assimilation

The mean carbon assimilation of dead *Halophila* (0.06± 0.32 SD *µ*mol C m^−2^ s^−1^) and *Thalassia* (0.12 ± 0.84 *µ*mol C m^−2^ s^−1^) were similar and remained relatively constant over the range of tested relative biomasses (Figure 2). In contrast, dead *Halodule* had higher variability (-0.1 ± 2.25 *µ*mol C m^−2^ s^−1^) and *Syringodium* rates were significantly higher (5.83 ± 2.97 *µ*mol C m^−2^ s^−1^) before steeply dropping to 0 at 33% relative biomass (Figure 2). All natural seagrass treatments show roughly linear trends with highest assimilation rates at 100% relative biomass reaching 0 carbon assimilation between 30 – 40% relative biomass (Figure 2). After correction (natural – dead = residual), *Halophila* had the highest maximum carbon assimilation rate (6.06 *µ*mol C m^−2^ s^−1^), followed by *Thalassia* (5.58 *µ*mol C m^−2^ s^−1^), *Halodule* (4.75 *µ*mol C m^−2^ s^−1^), and *Syringodium* (3.63 *µ*mol C m^−2^ s^−1^; Table 2). *Halophila, Halodule*, and *Thalassia* switched from positive carbon assimilation to negative (i.e., carbon release) at relative biomasses of less than 33% whereas *Syringodium* began to assimilate carbon.

**Table 2.**
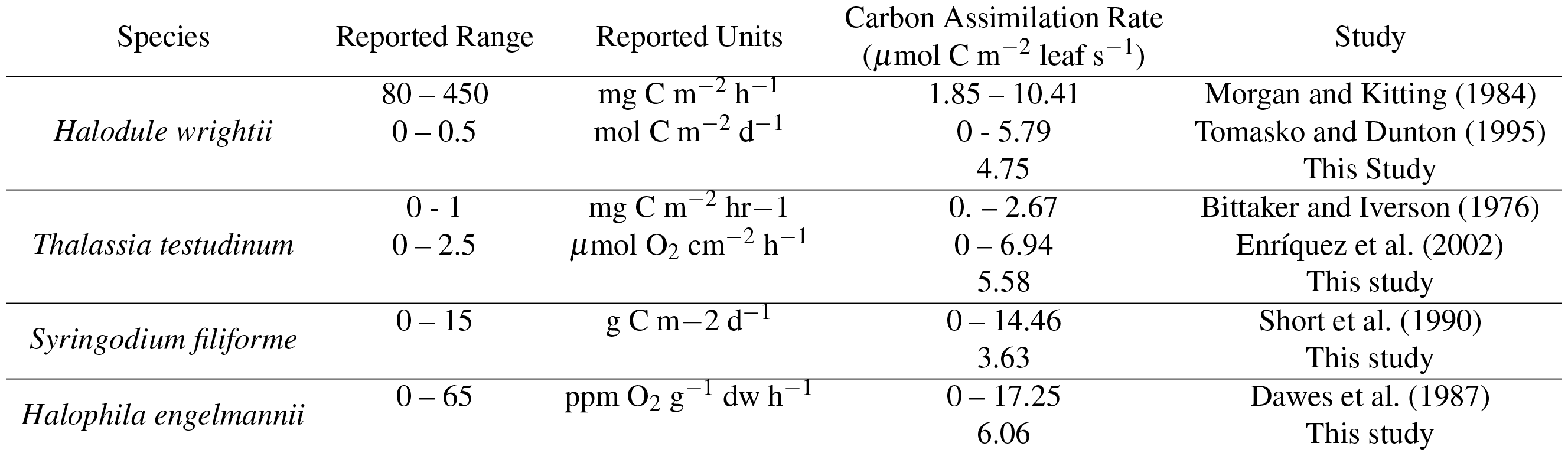
Comparison of carbon assimilation rates for Texas seagrasses.

**Figure 2.**
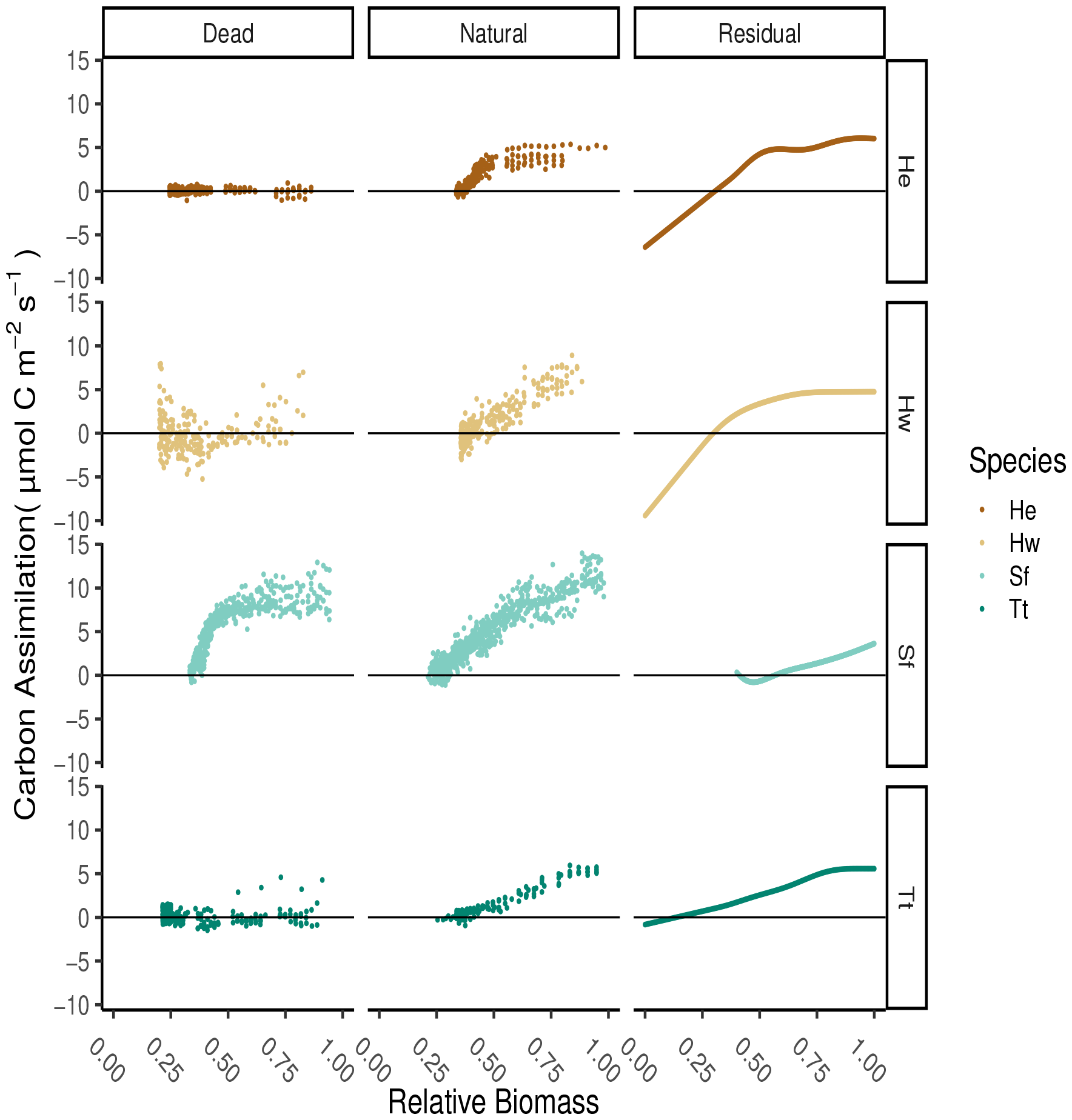
Dead seagrass blades produce species-specific carbon assimilation response curves, likely related to their morphology. Similar decreasing assimilation patterns are observed between all experimental blades. The residual model (experimental – dead) shows that carbon assimilation increases as relative biomass (i.e., water content) increases for all seagrass species. Note: the lower relative biomass of Sf is restricted because the strong exponential decay in the dead treatment would likely taper off.

### Effects of Desiccation on Maximum Leaf Quantum Efficiency

Before desiccation, the average maximum quantum efficiency of photosystem II (Fv/Fm) for all seagrass species was 0.79 ± 0.04 SD (Figure 3). However, the response of Fv/Fm to desiccation varied among species. The Fv/Fm dropped by *∼* 0.2 over 60 minutes for *Halodule* blades with no strong patterns between relative biomass and Fv/Fm (Figure 3). In contrast, all other seagrass species had ratio drops over 0.4 and showed strong relationships between Fv/Fm and relative biomass (Figure 3B). Overall, *Syringodium* had the highest biomass loss in this experiment and showed the strongest loss in Fv/Fm over time (Figure 3A). Some *Thalassia* did not follow this pattern and were able to maintain high Fv/Fm readings at low relative biomasses (0.75 at 25% biomass).

**Figure 3.**
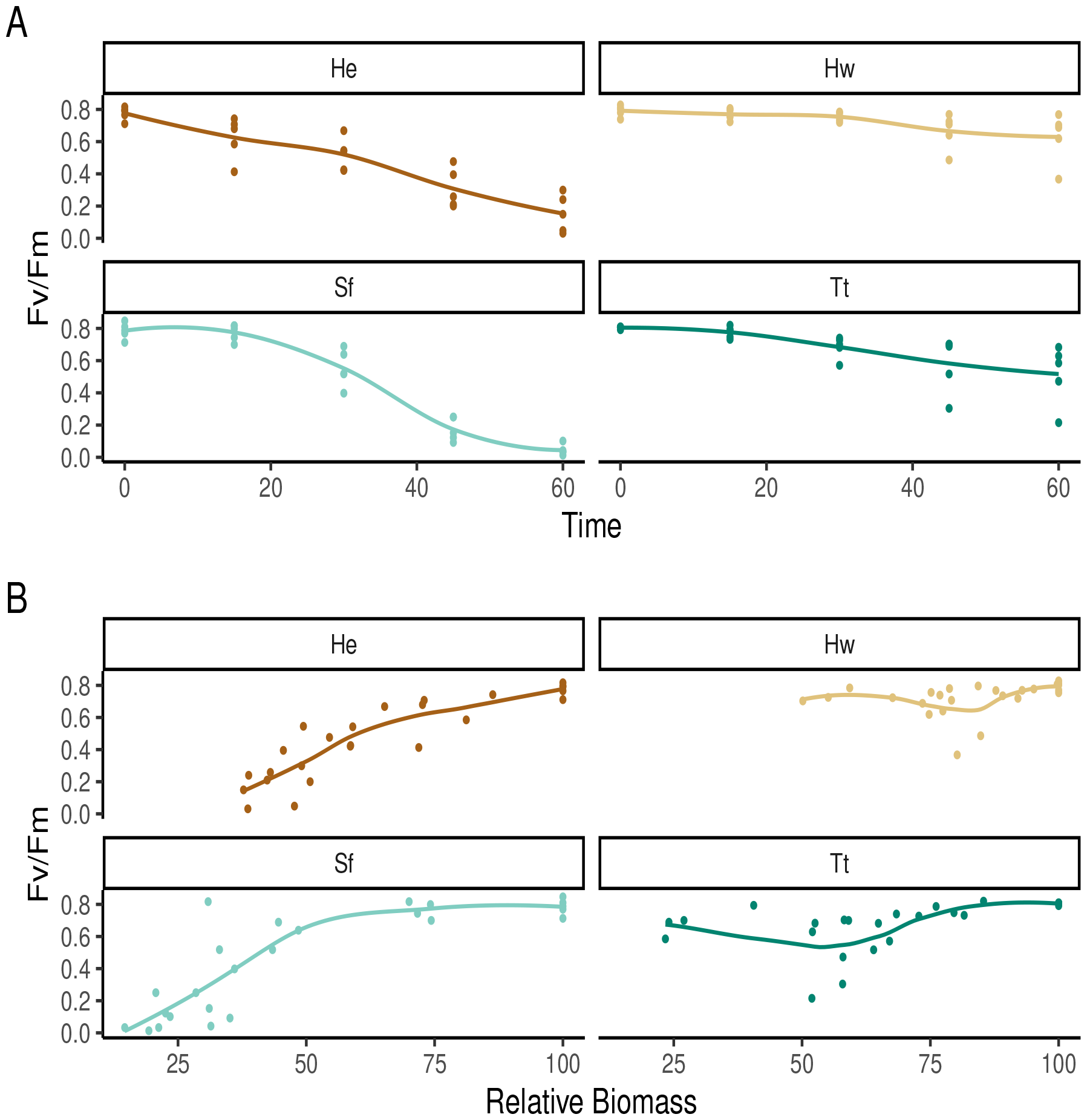
Changes in maximum leaf quantum efficiency over changes in A) time and B) relative biomass.

### Effects of Desiccation on Light Curves

The relative maximum electron transport rate (rETR_*max*_) of *Halophila* (32.2) was the highest of all tested seagrass species *Halodule* (18.5), *Syringodium* (14.7), and *Thalassia* (8.92; Figure 4; Table 3). *Halophila* needed the highest amount of irradiance for saturation (84.2 *µ*mol photons m^−2^ s^−1^) followed by *Halodule* (76.8), and *Syringodium* (41). The saturation irradiance of *Thalassia* from the LI-6400 (34.6) was similar to that measured by PAM (26.4). The initial photosynthetic rate was similar between *Halophila* (0.383), *Syringodium* (0.357), and *Thalassia* (0.338) but lower in *Halodule* (0.241). After 60 minutes of desiccation, only *Halodule* was able to create a light curve (Figure 4). All photosynthetic parameters were depressed compared to the non-desiccated curve (Table 3).

**Table 3.** Comparison of photosynthetic parameters from rapid light curves.

| Species | Instrument | Desiccation Time (min) | Maximum Photosynthetic Rate/Quantum Efficiency ($P_{max}/rETR_{max}$) | Minimum Saturating Irradiance ($E_k/ISat$) | Initial Photosynthetic Rate/Quantum Efficiency ($\alpha$) |
| --- | --- | --- | --- | --- | --- |
| *H. engelmannii* | PAM | 0 | 32.2 | 84.2 | 0.383 |
| *H. engelmannii* | PAM | 60 | - | - | - |
| *H. wrightii* | PAM | 0 | 18.5 | 76.8 | 0.241 |
| *H. wrightii* | PAM | 60 | 6.22 | 53.9 | 0.155 |
| *S. filiforme* | PAM | 0 | 14.7 | 41 | 0.357 |
| *S. filiforme* | PAM | 60 | - | - | - |
| *T. testudinum* | PAM | 0 | 8.92 | 26.4 | 0.338 |
| *T. testudinum* | PAM | 60 | - | - | - |
| *T. testudinum* | LI-6400 | 0 | 3.07 | 34.6 | 0.093 |

**Figure 4.**
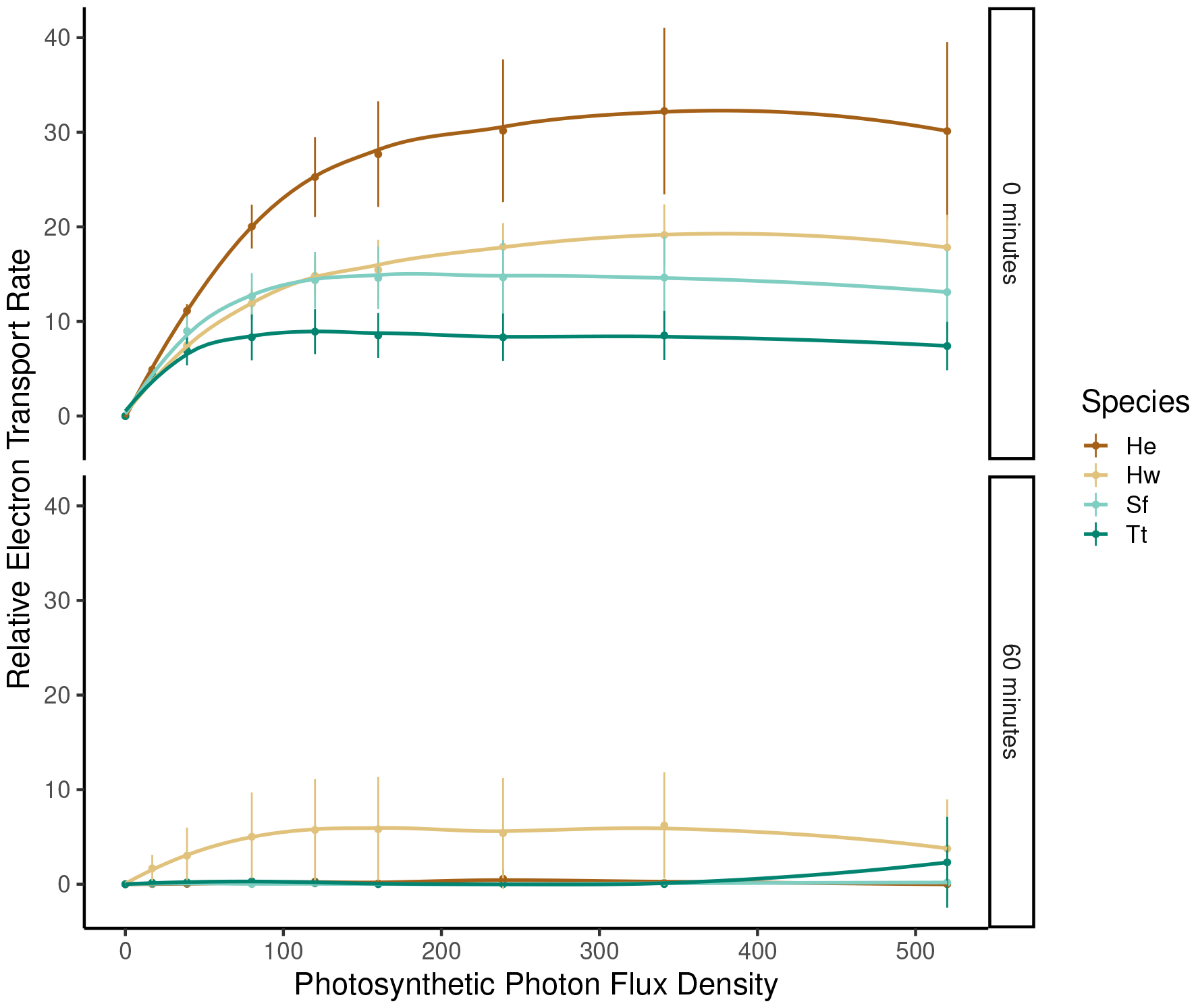
Rapid light curves at the initiation (0 minutes) and commencement (60 minutes) of desiccation stress. Each point represents a mean *±* standard deviation (n = 5).

### Timing and Frequency of Desiccation

The modeled average annual water evaporation rate for a seagrass meadow in south Texas is 0.84 ± 0.76 (SD) kg H_2_O m^−2^ h^−1^ with rates higher in summer (1.36 ± 0..71), lower in winter (0.41 ± 0.47) and fall (0.31 ± 0.46), and intermediate in spring (0.91 ± 0.75; Figure 5). Seagrasses spent longer exposed to the air in the winter (1807 hours) compared to the summer (1546 hours) with nearly no exposure during spring (440 hours) or fall (90 hours). Integrating over the season, more water would be lost during summer aerial exposures (8394 kg H_2_O m^−2^) than winter (2936 kg H_2_O m^−2^).

**Figure 5.**
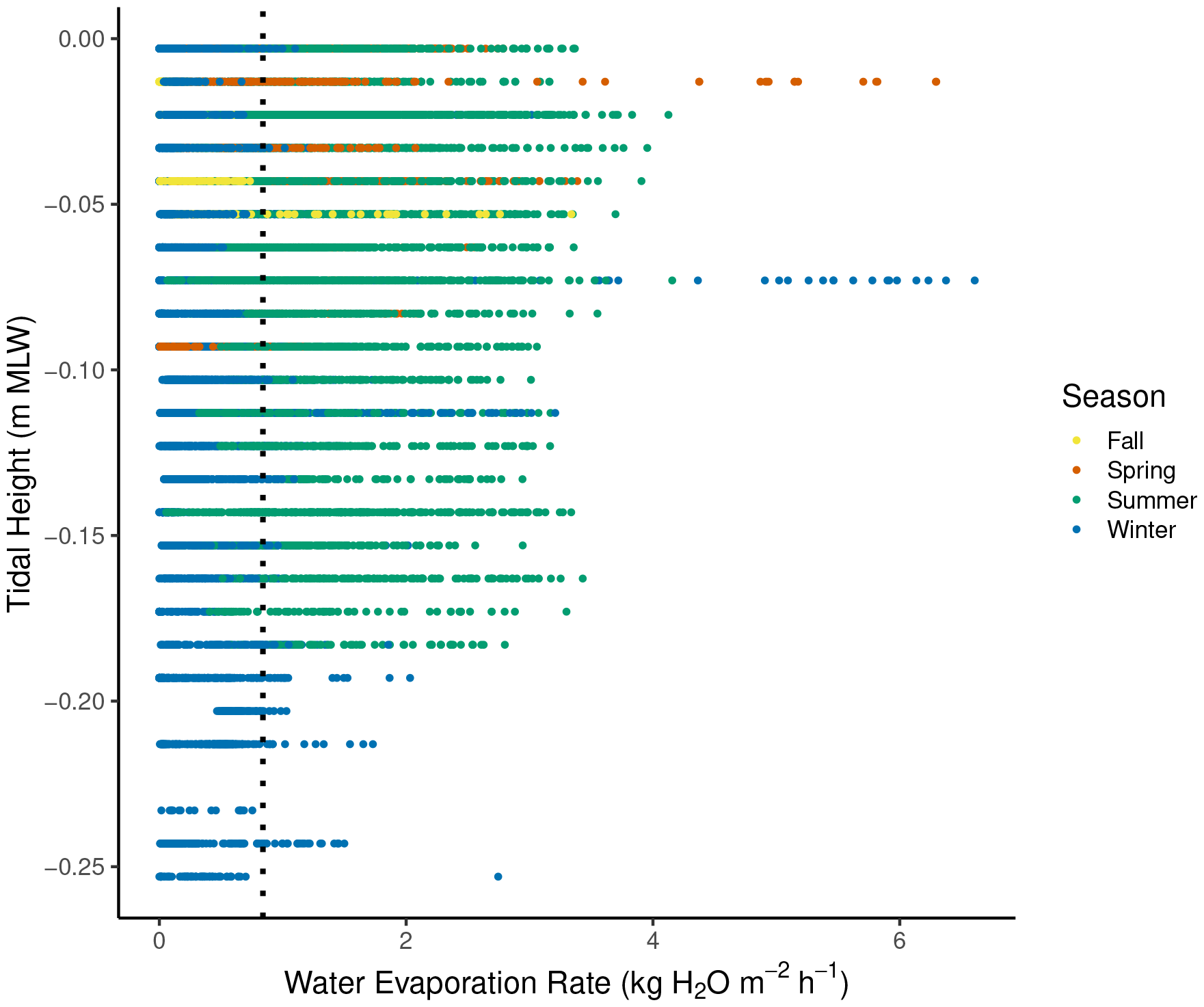
Modeled water evaporation using temperature, humidity, and wind speed for Port Aransas, TX from 2021 – 2023 during seagrass exposure. There are few spring and fall exposures because of Texas’ secular tides. The dashed vertical line represents the average water evaporation rate during this two-year period.

## DISCUSSION

This study assesses multiple techniques of measuring seagrass productivity to evaluate carbon assimilation potential under desiccation stress. Infrared gas analysis was applied to subtidal seagrass species to measure seagrass productivity through the lens of desiccation stress. Seagrasses did not show signs of stress (measured through pulse amplitude modulated“PAM” fluorometry) during measurements. We show that measurements fall within previously recorded values and align with expected ecological theory. We found that desiccation rates would be highest during the summer when other abiotic stressors are present. Overall, we recommend the usage of infrared gas analysis for seagrass productivity measurements.

### Occurrence and Timing of Desiccation Stress

Few seagrass species can grow within typical intertidal zones (e.g., *Zostera marina, Zostera noltii, Zostera japonica, Halophila ovalis, Thalassia hemprichii*) and are restricted to subtidal habitats (Den Hartog and Kuo, 2006). Subtidal seagrasses may become partially exposed to air when tides drop below the mean low water Koch (2001). Phillips (1960) noted water depths of 15 cm exposed rigid *Thalassia testudinum* to air causing sufficient desiccation stress to damage leaves. Even microtidal estuaries (e.g., Texas) have strong enough tidal excursions to expose populations of *Thalassia, Halod-ule*, and *Syringodium* because of low seasonal secular tides during summer and winter.

Previous work in Florida noted wintertime exposures were particularly stressful because of low humidity polar air and strong winds (Phillips, 1960; Strawn, 1961; Holmquist et al., 1989). In contrast, we show that most evaporation occurs during summertime exposure on the Texas Gulf Coast. Therefore, the timing of peak desiccation stress is location specific. During the Texas summer, seagrasses would experience compounding stressors (desiccation, light, salinity, thermal) that would likely decrease overall plant productivity and quickly lead to plant die offs.

### Seagrass Desiccation Tolerance

Abiotic and biotic factors shape common zonation patterns of many sessile marine organisms (e.g., seaweeds, invertebrates) controlling their upper and lower limits respectively Stephenson and Stephenson (1949). Numerous stresses affect these creatures (e.g., temperature, redox, mechanical), but desiccation remains one of the most intensely studied stress in plants. However, tropical seagrasses do not follow traditional paradigms because of their morphological and physiological adaptations. For example, Björk et al. (1999) found that seagrass species growing in the high intertidal zone possessed anatomical adaptations that minimized water loss during aerial exposure compared to less exposed species. Additionally, *Zostera japonica* (upper intertidal) is more sensitive to desiccation than *Zostera marina* (subtidal; lower intertidal) because of decreased photosynthetic responses at the leaf level (Shafer et al., 2007). Indeed, we found that desiccation rates decreased for deeper water species *Halodule wrightii* (30.17% min^−1^ cm^−2^), *Syringodium filiforme* (4.79% min^−1^ cm^−2^), *Thalassia testudinum* (2.3% min^−1^ cm^−2^). We were unable to test Hw stress levels at 25% relative biomass, but Tt remained unstressed and productive compared to Sf. Therefore, our data support previous studies showing that seagrasses show reverse desiccation tolerance patterns compared to algae.

### Carbon Fixation of Texas Seagrasses

Here, we report measurements of seagrass productivity directly referenced to the analyte of interest, carbon, over meaningful scales of variability. Each of our measured carbon assimilation rates fell within previously reported ranges for each species, indicating that our measurements are reasonable (Table 3). All plants were acclimated to the same environmental conditions, but there were species-level differences He *>* Tt *>* Hw *>* Sf. *Halophila* spp. are often understudied despite making up roughly a quarter of seagrass diversity (Dawes et al., 1987), but the few well studied examples (e.g., *H. ovalis*) are known to be highly productive (Erftemeijer and Stapel, 1999). The relationship between *Thalassia testudinum, Syringodium filiforme*, and *Halodule wrightii* is a well-known successional pattern where Hw is an early colonizer that is replaced by Sf, before endstage meadow of Tt (Williams, 1990). The environmental tolerances of Sf are narrower than Hw, likely explaining its decreased productivity at the tested conditions (CapistrantFossa and Dunton, 2024). Overall, the productivity measurements from the instrument reasonably match ecological theory and give direct rates of carbon assimilation.

### Evaluation of Li6400 as a Viable Strategy of Measurement of Carbon Fixation

Maximum leaf quantum efficiency (Fv/Fm) values are conserved between plant lineages and range from 0.7 – 0.83 for unstressed plants to *<* 0.7 for stressed plants (Maxwell and Johnson, 2000; Ritchie, 2006). Seagrasses were unstressed in our trials until they reached *∼*70% relative biomass. This is equivalent to *∼*1.5 minutes of desiccation in the LI-6400 chamber, more than the time required for instantaneous plant measurements. Further parameterization is needed to slow desiccation rates to produce accurate light curves because of the relatively long duration. Newer technologies (i.e., LI-6800) offer finer control over humidity in the chamber which would slow desiccation rate, or water can be added to the CO_2_ scrubber solution. We note that the calculated saturation irradiance of *Thalassia* was reasonably close between the PAM and LI-6400, indicating agreement between the two systems. Measurements of dead seagrass blades are needed to perform species-specific corrections to measurements. Here, we show little to no correction is needed for *Halophila* and *Thalassia* whereas *Halodule* and *Syringodium* did. We hypothesize the strong apparent assimilation of *Syringodium* and *Halodule* are because of their terete-like morphology with relatively large internal volumes that can hold larger internal pressures of CO_2_ than the flat, thin, blades of *Thalassia* and *Halophila*.

## CONCLUSIONS

Previous studies have shown that IRGA can successfully be used to measure carbon assimilation in intertidal seagrass species because of their relative ease of access (e.g., Leuschner and Rees, 1993; Leuschner et al., 1998; Silva et al., 2005). However, subtidal seagrasses can also be easily accessed because can be temporarily exposed by low tides, have a longer blade length than water column depth, or be dug from the sediment. Given that plants appear to be relatively unstressed during the short timescales of emersion we recommend that this technology be further investigated for use in the field for non-invasive to minimally destructive sampling of subtidal populations seagrass.

We note major benefits include: 1) the direct measurement of carbon assimilation rather than oxygen evolution removing the need for a highly variable photosynthetic quotient thereby improving productivity measures, 2) improving accuracy of desiccation studies by removing the need for immersion into liquid media for productivity measurements after drying (e.g., Dring, 1982), 3) rapidly evolving technology for underwater-based productivity measurements, and 4) allowing for ecologically relevant differentiation of productivity between aerial exposure and inundation for accurate carbon budgeting.

## ACKNOWLEDGMENTS

We sincerely thank J. Reid Fischer for his assistance in the initial planning, testing, and fieldwork with the LI-6400 XT and Walz Diving PAM experiments. Additionally, we would like to thank Lisa Young and Anastasia Canu for field and lab support needed to make this project successful.

